# Modeling predicts that CRISPR-based activators, unlike CRISPR-based repressors, scale well with increasing gRNA competition and dCas9 bottlenecking

**DOI:** 10.1101/719278

**Authors:** Samuel Clamons, Richard Murray

## Abstract

Synthetic transcriptional networks built from CRISPR-based repressors (CRISPRi) rely on shared use of a core dCas9 protein. In *E. coli*, CRISPRi cannot support more than about a dozen simultaneous gRNAs before the fold repression of any individual gRNA drops below 10x. We show with a simple model based on previous characterization of competition in CRISPRi that activation by CRISPR-based activators (CRISPRa) is much less sensitive to dCas9 bottle-necking than CRISPRi. We predict that *E. coli* should be able to support dozens to hundreds of CRISPRa gRNAs at >10-fold activation.

## 1. Introduction

One of the most powerful and flexible tools in the modern synthetic biologist’s toolkit is the synthetic transcription factor network, which computes and actuates using cascades of activation or repression by either naturally-occurring or engineered transcription factors. Synthetic variants of transcription factor networks have been used to build molecular oscillators [1, 2, 3, 4], molecular fold change detectors [5, 6], signal level discriminators [7], and simple, composable elementary logic units for both analog [7] and digital [8, 9, 10, 11] computation.

To date, however, synthetic transcription factor networks remain limited in size, with the largest circuits containing on order of a dozen transcription factors [9, 10]. Several constraints limit the size of these networks. We focus on two such constraints here:

- **Difficulties of part design:** Put simply, transcription factors are hard to make. Traditional, wild-mined transcription factors like TetR, LacI, LasR, AraC, etc., are invaluable for small-scale prototyping, but it is difficult to find large sets of transcription factors with compatible operating concentrations and no crosstalk (the largest known set currently consists of the 12 mutually-orthogonal transcription factors in [9]). In principle, an almost arbitrary number of transcription factors could be built using zinc finger nuclease (ZFN) or transcription activator-like effector nuclease (TALEN) technology, but in practice, high-quality ZFNs are difficult to engineer and TALENs are prone to recombination and mutational breakage.
- **Difficulties of limited resources:** Transcription factor networks use up cell resources (ATP, amino acid, ribosomes, etc.). Particularly in bacteria and other small, resource-limited cells, this load can cause unexpected “retroactivity” feedback between nominally-unconnected circuit components [12, 13]. Resource limits are particularly problematic when they interfere with host cell growth, leading to evolutionary pressure to disable the offending circuit [14, 15].

One promising development in synthetic transcription factor design is the use of mutationally-inactivated Cas9 programmable nucleases [16] as either repressors (CRISPRi; [16, 17, 18, 10, 19]) or activators (CRISPRa; [18, 20, 21, 22, 23]). The simplest example of a CRISPR transcription factor (CRISPRtf) is dCas9, which is Cas9 with two mutations that disable its ability to cut DNA. When targeted to a promoter by a guide RNA (gRNA), dCas9 strongly binds to the promoter, blocking RNA polymerase attachment or transcriptional elongation (at least, in prokaryotes). Other CRISPRtfs have been made by either fusing native polymerase-recruiting factors to dCas9 or by recruiting those factors to a binding domain on the gRNA. These synthetic transcription factors can be easily modified to create new, orthogonal versions by simply changing the sequences of the binding site and gRNA.

CRISPRtfs are easy to generate and have been used to successfully manipulate host gene expression [8] and to build limited-scale synthetic transcriptional circuits [10]. However, while CRISPRtfs largely bypass difficulties of part design, they introduce a new barrier to scalability in the form of a troublesome resource bottleneck – dCas9 or dCas9 fusion protein. Any expression of one gRNA in a network of CRISPRtfs sequesters shared dCas9 away from other gRNAs, decreasing their effectiveness as outlined in detail in [24]. The dCas9 bottleneck can be loosened by producing more dCas9, but this strategy is limited by the toxicity of dCas9 when expressed at high concentration, especially in prokaryotes [25, 26].

In [27], Zhang and Voigt quantify this bottlenecking effect in *E. coli* They show that under physiological, circuit-like conditions, dCas9 bottlenecking causes CRISPRi target repression to drop off as roughly 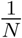, where *N* is the number of competing gRNAs expressed. Furthermore, they show that *E. coli* expressing near-maximal sustainable levels of dCas9 repressor cannot support more than about ∼ 7 simultaneous gRNAs at a 10-fold level of repression (or about ∼ 15 simultaneous gRNAs using a low-toxicity dCas9 variant, dCas9*-PhlF).

Prior to 2018, CRISPRtfs in bacteria were largely limited to repressors, as existing CRISPR activators typically exhibited ≤ 10-fold activation and only functioned in an unusual, *RpoZ*-knockout strain; accordingly, the Zhang and Voigt’s analysis of resource bottlenecking reasonably considered only CRISPRi. There are now more effective CRISPRa activators made from *SoxS* fusion proteins that do not require *RpoZ* knockout, which makes CRISPRa a feasible alternative to CRISPRi for synthetic gene circuit construction [23].

In this paper, we use a simple model of gRNA competition for dCas9 to show that CRISPR activators are substantially less vulnerable than CRISPR repressors to dCas9 bottlenecking. Our model anticipates that CRISPRa should be able to support many times more simultaneous gRNAs than CRISPRi under most conditions, although CRISPRi may be more effective for very small networks under ideal conditions.

## 2. Why activators are less sensitive to bottlenecking than repressors

CRISPR activators should scale better than CRISPR repressors because, in general, activators are more robust than repressors against removal of regulator when they are already saturating.

Consider a repressor and an activator of equal strength and at high enough concentration that every target promoter is bound. What happens if the concentration of each regulator drops enough that one of the targets becomes unbound? How much is each system affected? We could equivalently consider the fraction of time bound for a single-target system, but for conceptual simplicity, we will consider a system with a “large number” (say, > 5) of targets that are each either bound or not.

When one activator drops off a target, total expression falls by a little less than a fraction 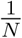 of maximum expression, where *N* is the number of target promoters. The fold change in expression caused in this decrease in activators is therefore roughly 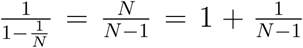. If *N* is reasonably large, this fold change will be quite small (consider *N* = 10).

When all target promoters are bound by repressors, the total expression of the bound promoters is 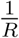, where *R* is the fold-repression caused by a single repressor binding to a single promoter. When a single repressor is removed, the change (increase, this time) in expression is, again, roughly 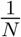, but this change occurs against a background of “leak” experienced when all repressors are bound (which is hopefully a small value) rather than maximum possible promoter expression (which is hopefully a larger value). The fold change experienced due to this removal is roughly 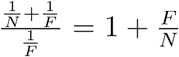. This fold change is only small if the number of targets is large compared to the fold-repression of the repressor, and for most reasonably capable repressors in most reasonably-sized cells it should be at least 2-fold.

Viewed another way, the changes in expression experienced when a small amount of activator or repressor are removed are of roughly the same *absolute* magnitude, but the relative impacts of those effect are quite different – for an activator, the change should be compared against maximum expression (which should be large), whereas for a repressor, the change should be compared against promoter leak (which should be small). Figure 1 shows this difference for a concrete example of a 10-fold activator and a 10-fold repressor targeting a 5-copy promoter.

**Figure 1:**
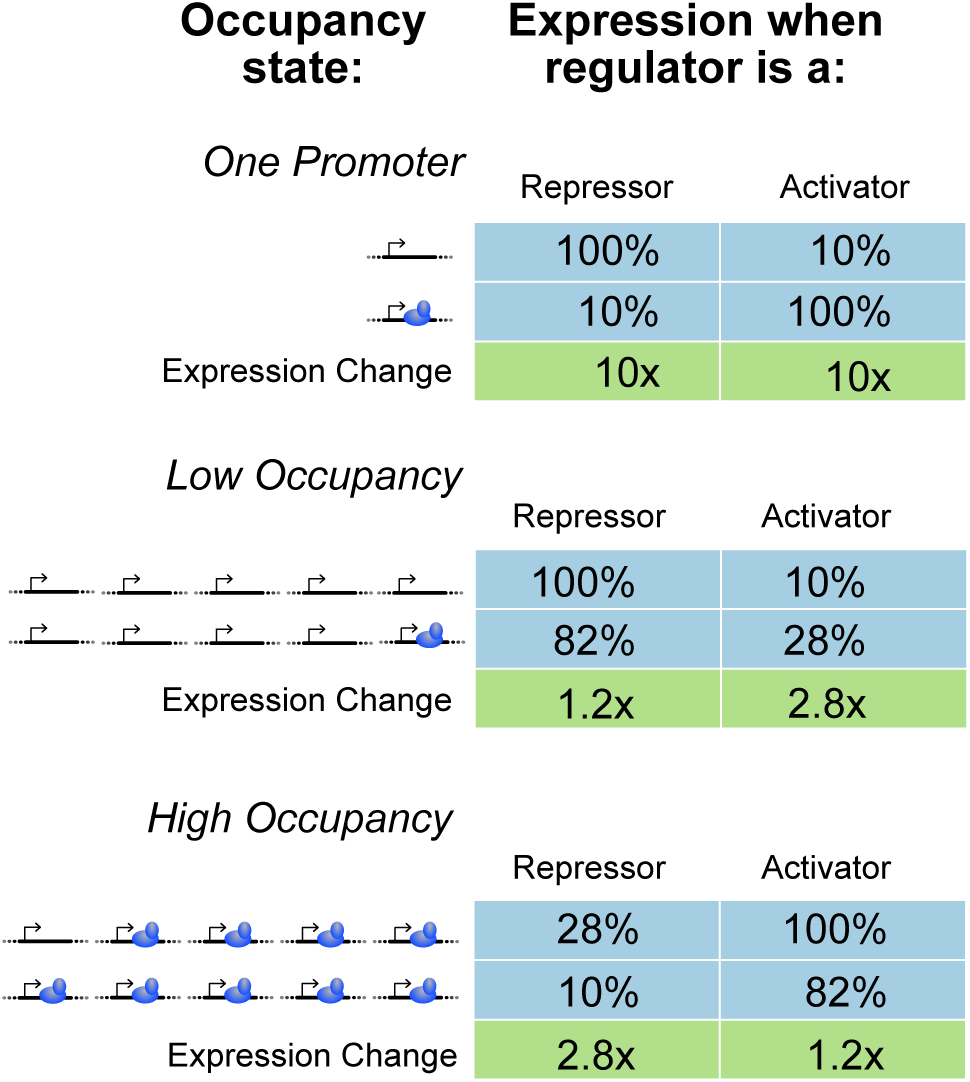
Changes in expression when the concentrations of two regulators (an activator and a repressor) are changed under different regulator concentration regimes. Both the repressor and the activator change expression 10-fold (top table). At low saturation of target promoter by the regulators, the activated promoter is more sensitive to changes in regulator concentration than the regulated promoter (middle table). Conversely, at high target saturation, the activated promoter is more robust to changes in regulator concentration (bottom table).

Note that the asymmetry between sensitivities of activators and repressors reverses itself at low concentrations of regulator – there, activators are more sensitive to regulator changes and repressors are more robust. Compare Figure 1, middle and bottom rows. Activators trade off high sensitivity to concentration changes at low saturation for robustness to concentration changes at high saturation, while repressors make the opposite tradeoff.

## 3. Simulations show that CRISPRa scales better than CRISPRi

Using a steady-state solution for a simple ODE model of CRISPR, we numerically investigated the effects of gRNA competition on minimal CRISPRi and CRISPRa systems consisting of a repressing or activating dCas9:gRNA complex targeting a reporter gene. See section 5 for a description of the model and details on how we calculate fold change.

We calculate fold change for three different concentrations of target promoter roughly representing a genomically-integrated reporter, a reporter on a low-copy plasmid, and a reporter on a high-copy plasmid (Fig. 2A, B, and C, respectively). We use parameters estimated for this model from *in vivo* data by Zhang & Voigt ([27]). We vary the concentration of dCas9 from 100 nM up to 530 nM, which is roughly the maximum concentration of dCas9 the E. coli used by Zhang & Voigt can support before suffering significant growth defects.

**Figure 2:**
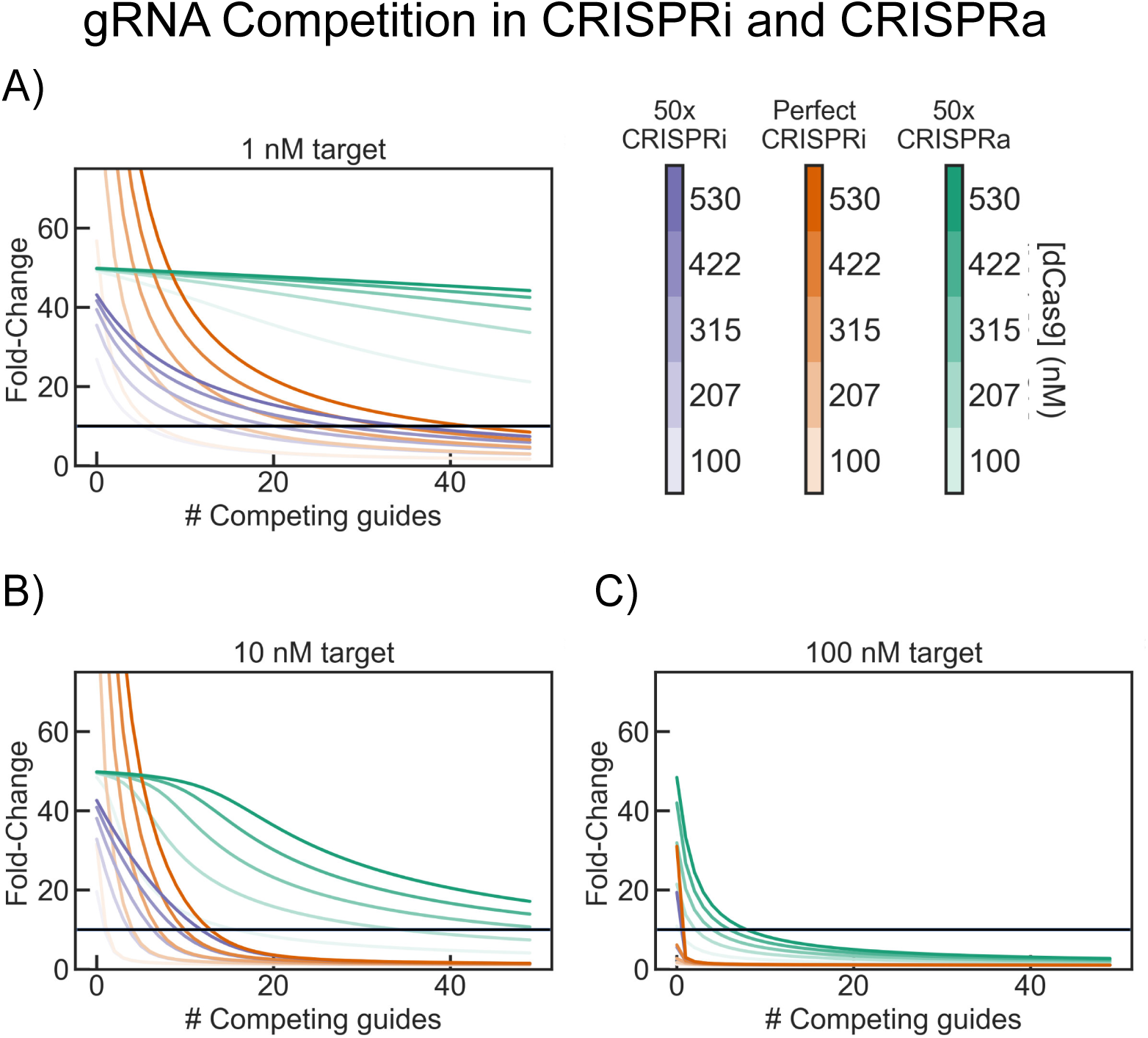
Simulated fold change of activation with CRISPRa (green) or repression with CRISPRi (perfect repression, orange; or 50x repression, purple) with various dCas9 concentrations and either A) 1 nM, B) 10 nM, or C) 100 nM of target promoter. Parameters taken from [27]. The horizontal line in each plot marks the10x fold change.

The same model that (correctly) predicts that the scalability of CRISPRi is severely hampered by inter-gRNA competition also predicts that CRISPRa should be far more robust against inter-gRNA competition.

We also predict that CRISPRi may produce higher-fold changes than CRISPRa at low numbers of competing guide RNAs, especially when dCas9 is abundant with respect to the target promoter. Note that this prediction is a direct consequence of the assumption that dCas9 is a perfect repressor, but a finite activator. This means that the effectiveness of CRISPRi repression is unbounded above – as binding becomes more efficient, the fold change of repression approaches infinity – while CRISPRa activation effectiveness is bounded above at 50x. If we relax this assumption so that CRISPRi is equally as “effective” as CRISPRa (i.e., it represses 50x when fully bound), then it fails to exceed (or even match) the overall fold-repression of CRISPRa at *any* number of competing gRNAs (Fig. 2, purple curves).

These results are not unduly sensitive to the particular parameters estimated in [27]. We computed maximum “acceptable” competing gRNA number (the largest number of competing gRNAs that still allows 10 fold regulation of the target) for 1,000 randomly sampled parameters (see Fig. 4 for results from 100 representative parameter sets; see figure legend for details on parameter sampling). Although there was a significant degree of variation in CRISPRa scalability across simulations, only rarely did we observe CRISPRa with worse scaling than any simulated CRISPRi system (Fig. 4B).

**Figure 3:**
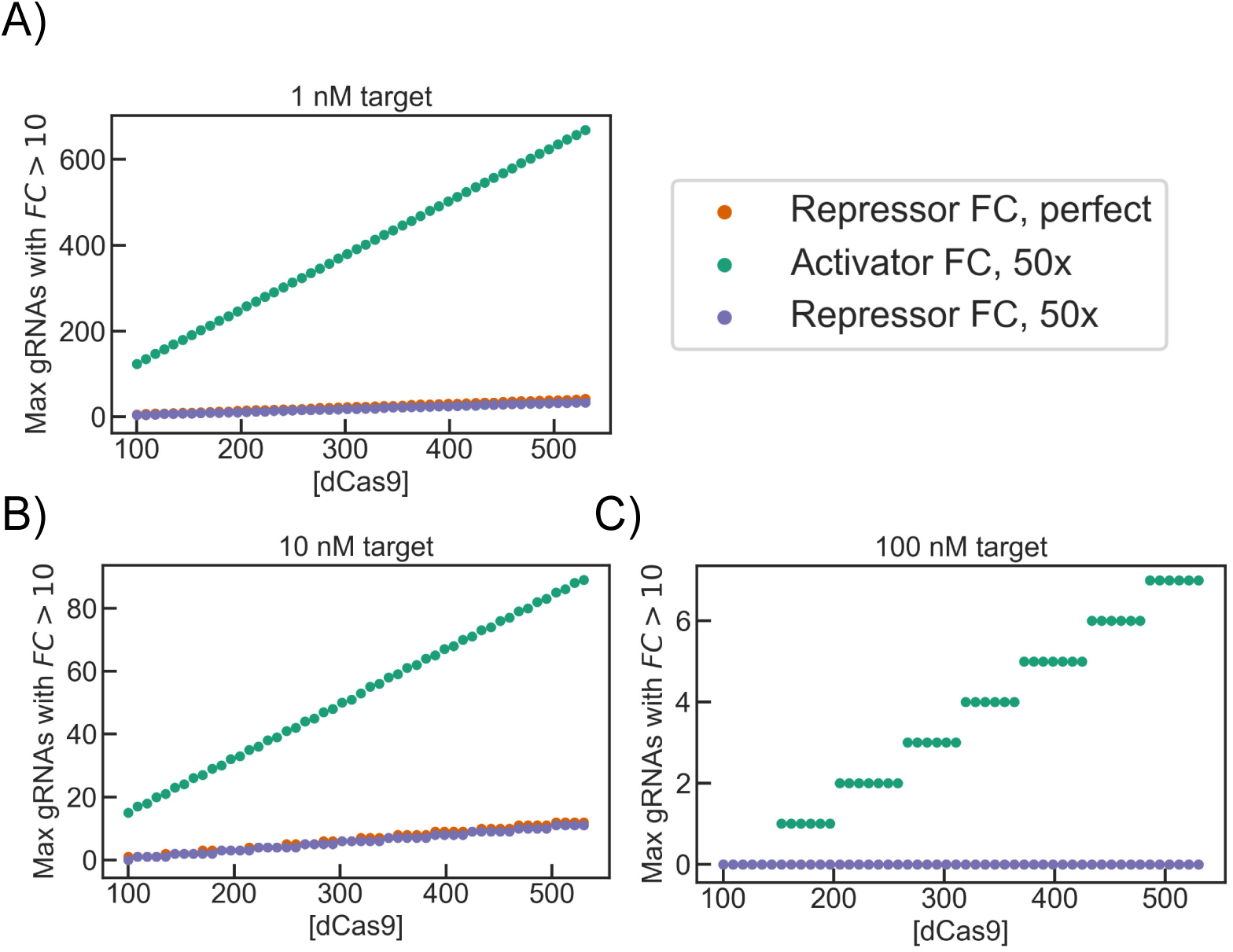
Predicted maximum number of competing guide RNAs before competition reduces fold change to below 10x, for various dCas9 concentrations and A) 1 nM, B) 10 nM, or C) 100 nM of target promoter. Parameters taken from [27]. “0 Max gRNA” means that a single gRNA competitor is sufficient to drive fold change below 10x.

**Figure 4:**
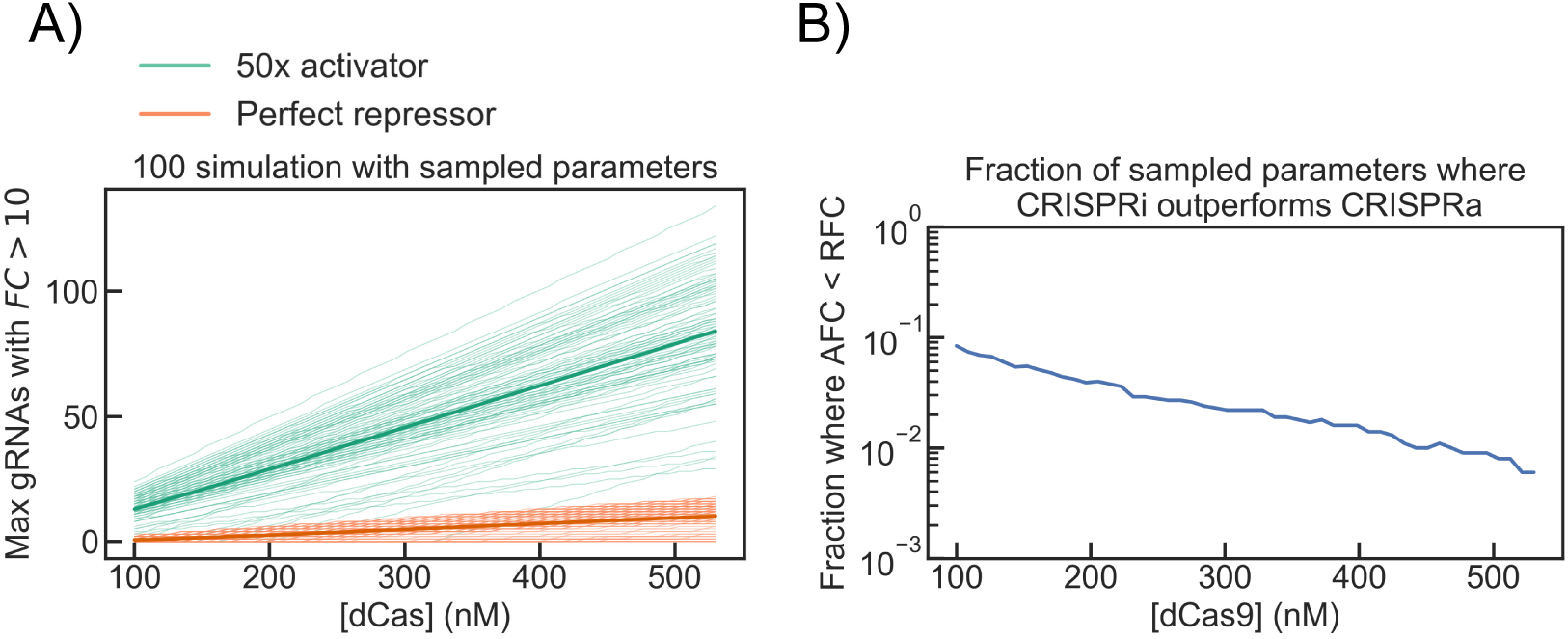
A) Predicted maximum number of competing guide RNAs that still allow 10-fold regulation for 100 of the 1,000 tested CRISPRi/a systems with randomly sampled parameters. Thin lines represent single sampled parameter sets; bold lines represent averages across all sampled parameter sets. Target copy number was held fixed at 10. Transcription speed for competing gRNAs was held to a constant multiple of that of the primary gRNA (changes in one transcription rate relative to another are equivalent to a compression or expansion of the gRNA concentration axis). Fold-activation of the CRISPRa activator was sampled from a normal distribution with mean 50x and a 20% standard deviation, reflecting our high confidence in our estimates of that parameter from [23]. All other parameters were log-normally distributed around their best-estimate values, as determined in [27], with a log10 standard deviation of 0.5, very roughly representing a half-order-of-magnitude “50% confidence range”. B) The fraction of sampled parameter sets (out of 1,000) in which CRISPRa was predicted to perform better than *any* sampled CRISPRi, at each possible dCas9 concentration (CRISPRi was never observed to perform better than CRISPRa with the same parameters).

We also performed local gradient-based sensitivity analysis on each parameter. See Section 5.3 for details.

## 4. Discussion

There are good reasons to use CRISPRi instead of CRISPRa when building bacterial biocircuits, at least in the near future. CRISPRi requires fewer moving parts, produces larger best-case fold changes (and unquestionably higher fold changes in single-gRNA systems), and is less likely to drastically interfere with host genetic expression, at least in *E. coli*, as current-generation prokaryotic CRISPRa activators use modified versions of *E. coli* activators that upregulate some native promoters.

Repression is arguably also a more useful tool than activation. In particular, several simple, classic genetic circuits rely exclusively on repression (e.g., the repressilator [1], the two-gene genetic toggle switch [30], NOR-gates [10]).

Finally, CRISPRa is more difficult to use on natural genomic targets, as it requires a PAM within a ∼10 bp window of an ideal position upstream of the target promoter’s −35 box. For some genes, this target simply does not exist.

Nevertheless, CRISPRa has a distinct advantage over CRISPRi – it should be significantly less impacted by bottlenecking of core dCas9. CRISPRi effectiveness is predicted to drop precipitously for systems above about a dozen gRNAs, while CRISPRa should still function in the presence of dozens-to-hundreds of competing gRNAs.

As CRISPR-based synthetic circuits grow in scale and complexity past about a dozen components, we anticipate that the usefulness of CRISPRi will drop off precipitously, while CRISPRa should still function even in the presence of dozens of competing gRNAs (at least, for circuits with low target concentrations). We urge anyone who dreams of building large CRISPR-based biocircuits to consider using CRISPRa.

## 5. The Model

We adapt the modeling framework derived in [24] and adapted by Zhang and Voigt in their analysis of gRNA competition in CRISPRi. We recapitulate their analysis and extend it to CRISPRa.

### 5.1. A simple model of CRISPRi

Following the analysis in [27] (originally formulated by Chen, Qian, and Del Vecchio in [24]), we will consider a model of dCas9 repression in which a gRNA g1 is in competition with some number *N* of functionally identical, non-targeting gRNAs for a fixed amount of core dCas9. We wish to calculate the fold change of repression of the target of g1 as a function of *N*.

We model dCas9 as a Shea-Ackers repressor of cooperativity 1 and a single binding site per promoter [28]. We can then write down the average transcription rate of a target promoter as a sum of rates *r*_*f*_ and *r*_*C*_ of transcription from free and dCas9-bound promoter, respectively, weighted by the concentration of promoter in each of those states:

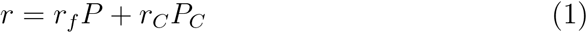

where *r* is the average transcription rate from the target promoter. If no gRNA is expressed, *P*_*C*_ = 0 and equation 1 simplifies to *r* = *r*_*f*_ *P*_*tot*_. We assume that *r*_*C*_ = 0 (i.e., repression by bound dCas9 is perfect, allowing no leak), so when the gRNA is active, equation 1 simplifies to *r*_*f*_ *P*, where *P* will vary with dCas9 concentration, gRNA expression level, number of competing gRNAs, etc. We can then write the fold change of repression (RFC) as

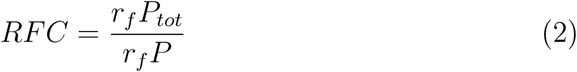

Now we can solve RFC at steady-state. We assume total promoter concentration is held constant, so *P*_*tot*_ = *P* + *P*_*C*_. At steady state, there will be flux balance between dCas9 binding to *P* and dCas9 unbinding from *P*_*C*_, so *P*_*C*_ = *KC*_*g*1_*P*, where 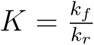 is the association constant of binding between dCas9:gRNA complex and free promoter. RFC then becomes

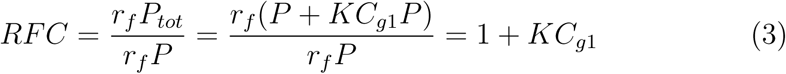

Now we make the informal quasi-steady state assumption that total dCas9 concentration *C*_*tot*_ is a constant set upstream by dCas9 production rates and dilution/degradation. Using a conservation law for dCas9 and steady-state flux balance of dCas9 and guide RNAs, the steady-state concentration of dCas9:g1 (*C*_*s*_1) can be calculated as:

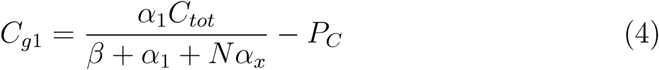

To find the steady-state concentration of *P*_*C*_, we use 1) flux-balance between bound and free promoter and 2) mass conservation of promoter:

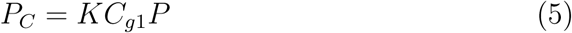

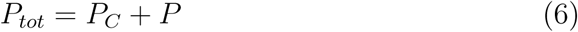

Combining these and rearranging to solve for *P*_*C*_, we get

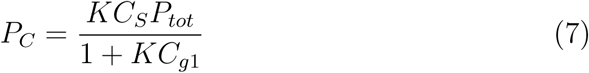

where, again, *K* is the *association* constant for binding between dCas9:gRNA and promoter. Substituting this into (4) and rearranging yields a quadratic polynomial in *C*_*S*_:

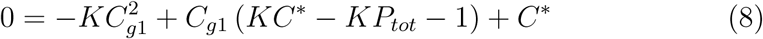

Where

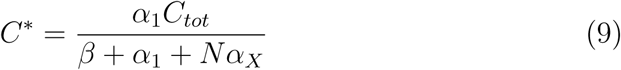

*C*_*g*1_ is then well-specified algebraically, and can easily be computed numerically.

### 5.2. A simple model of CRISPRa

We apply a similar analysis to the case of an activating CRISPR system under varying gRNA competition. The most salient changes from the analysis in section 5.1 are 1) *r*_*C*_ ≉ 0 (and *r*_*f*_ ≉ 0), and 2) the fold change we wish to calculate is the reciprocal of that in the CRISPRi case. This gives a fold change of activation (AFC) of

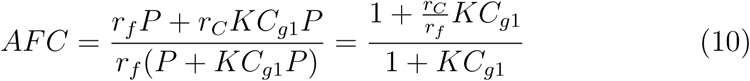

again with *C*_*g*1_ specified by Eq. (8) and (9). We are left with an additional parameter in the activation case, the ratio of activation 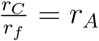 for a single target promoter by a bound activator. This is because we cannot assume zero expression from an un-activated promoter the way we assumed perfect repression of the bound promoter in CRISPRi – in practice, engineered activatable CRISPRa promoters are built from weak, but functional, constitutive core promoters that produce some transcription when unbound. We estimate an activation ratio of 50:1 based on the optimized dCas9-SoxS_*R*93*A*_ activation system from [23].

### 5.3. Sensitivity to Parameters

We investigated the robustness of our results to errors in parameters by subjecting our model to a local, gradient-based sensitivity analysis, using two summary statistics of CRISPRa performance relative to CRISPRi. We define the “fold change overperformance” of a pair of CRISPRa/CRISPRi systems with a specific set of parameters (including a specific number of competing guides) as the fold change of activation for a CRISPRa system with that set of parameters divided by the fold change of repression for a CRISPRi system with that set of parameters.

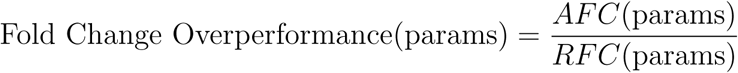

We also measure the advantage in scalability of CRISPRa over a CRISPRi system with the same parameters with a measure we call “scaling overperformance”, which we define as the maximum number of competing gRNAs that the system can tolerate with fold change caused by g1 remaining above 10-fold for the CRISPRa system, divided by the same number for the CRISPRi system.

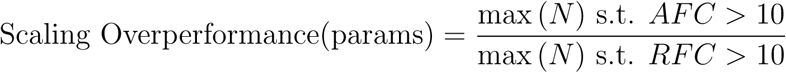

For each overperformance measure, we numerically calculate sensitivity of that measure to each parameter as the derivative of overperformance with respect to that parameter at our best-guess parameter set. These raw sensitivity values were normalized against parameter scale (i.e., errors in parameter estimation are assumed to be proportional in scale to the values of those parameters) and overperformance at the best-guess parameter value (so that sensitivity is given as a relative error).

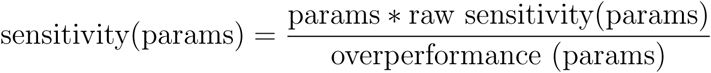

For *N* = 30 competing gRNAs, *P*_*tot*_ = 10 target copies, *C*_*tot*_ = 530 copies of dCas9 (roughly the maximum number an *E. coli* cell can support with negligible growth defect), *r*_*A*_ = 50, and all other parameters set as in Table 1, we find the sensitivities of fold change overperformance and scaling overperformance listed in Tables 3 and 4, respectively.

**Table 1:**
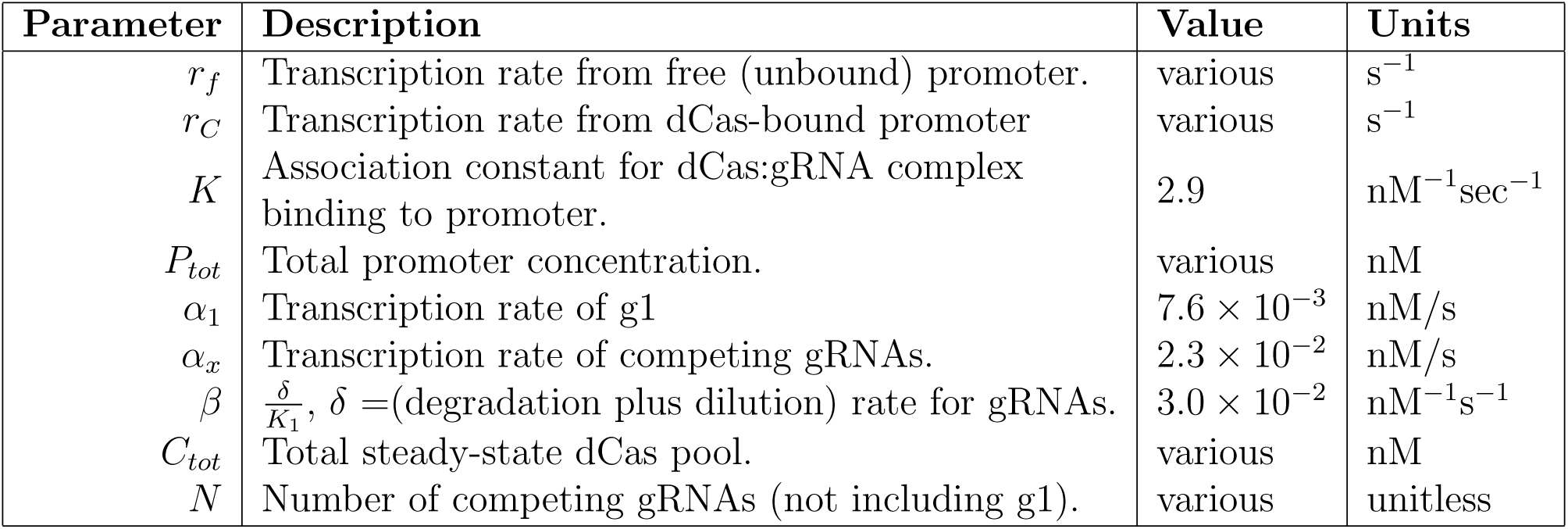
Parameter notation and values used in this model. Where possible, values were taken from fits to endpoint repression data in [27] (see Fig. 3 of that paper).

| Parameter | Description | Value | Units |
| --- | --- | --- | --- |
| $r_f$ | Transcription rate from free (unbound) promoter. | various | $s^{-1}$ |
| $r_C$ | Transcription rate from dCas-bound promoter | various | $s^{-1}$ |
| $K$ | Association constant for dCas:gRNA complex binding to promoter. | 2.9 | $nM^{-1}sec^{-1}$ |
| $P_{tot}$ | Total promoter concentration. | various | nM |
| $\alpha_1$ | Transcription rate of g1 | $7.6 \times 10^{-3}$ | nM/s |
| $\alpha_x$ | Transcription rate of competing gRNAs. | $2.3 \times 10^{-2}$ | nM/s |
| $\beta$ | $\frac{\delta}{K_1}$, $\delta$ =(degradation plus dilution) rate for gRNAs. | $3.0 \times 10^{-2}$ | $nM^{-1}s^{-1}$ |
| $C_{tot}$ | Total steady-state dCas pool. | various | nM |
| $N$ | Number of competing gRNAs (not including g1). | various | unitless |

**Table 2:** Dynamic variables used in our model.

| Variable | Description |
| --- | --- |
| $P$ | Concentration of free promoter. |
| $P_C$ | Concentration of promoter bound to dCas9 repressor or activator. |
| $C_{g1}$ | Concentration of dCas9:g1 complex. |
| $C$ | Concentration of free dCas9 regulator. |

**Table 3:**
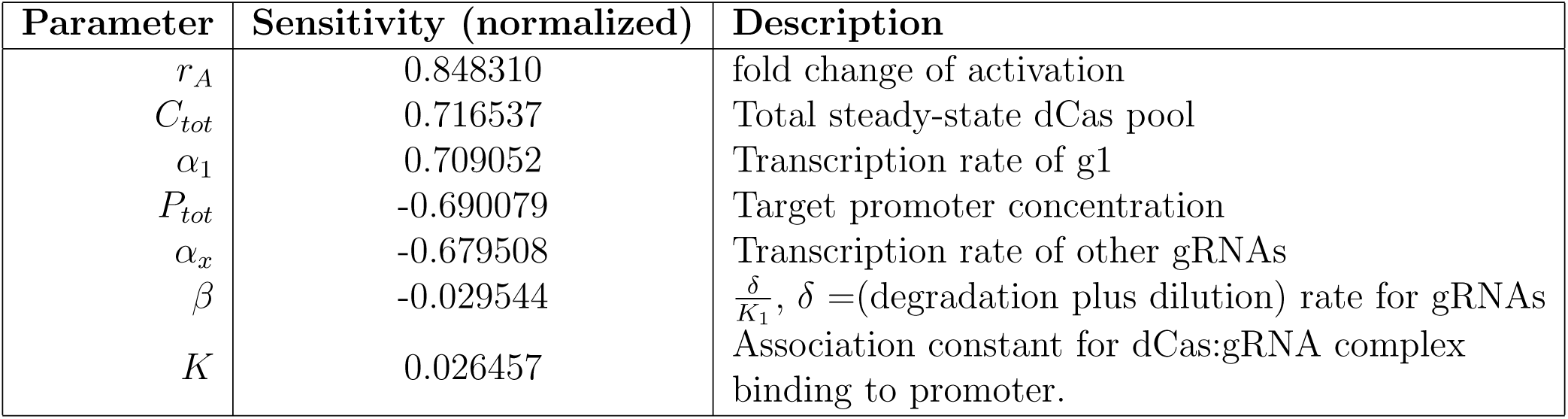
Sensitivity of fold change overperformance to each parameter.

**Table 4:** Sensitivity of scaling overperformance to each parameter.

| Parameter | Sensitivity (normalized) | Description |
| --- | --- | --- |
| $r_A$ | 1.079925 | fold change of activation |
| $C_{tot}$ | -0.335800 | Total steady-state dCas pool |
| $\alpha_1$ | -0.267926 | Transcription rate of g1 |
| $\beta$ | 0.267926 | $\frac{\delta}{K_1}$, $\delta$=(degradation plus dilution) rate for gRNAs |
| $P_{tot}$ | 0.207065 | Target promoter concentration |
| $K$ | -0.128736 | Association constant for dCas:gRNA complex binding to promoter. |
| $\alpha_x$ | -0.000000 | Transcription rate of other gRNAs |

Unsurprisingly, fold change overperformance is most sensitive to *r*_*A*_, the fold change of activation for promoters bound to a CRISPR activator. Fold change overperformance is also somewhat sensitive to *C*_*tot*_, *P*_*tot*_, *α*_1_, and *α*_*x*_. We have already shown how CRISPRi and CRISPRa performances change over realistic values of *C*_*tot*_ and *P*_*tot*_.

The parameters *α*_1_ and *α*_*x*_ – the transcriptional speeds of the target gRNA and competing gRNAs, respectively – are effectively different ways of scaling *N*. Increasing *α*_*x*_ changes the concentration of competing guide RNA in the same way that increasing *N* does, while changing *α*_1_ is equivalent to changing both *α*_*x*_ and *C*_*tot*_. Therefore, changes to these two variables are roughly equivalent to shifting the performance curves shown in Figure 1 along the “# Competing Guides” axis, albeit in a nonlinear way.

Fold change overperformance is relatively unaffected by *K* and *β*, which both incorporate binding constants and are therefore parameters of particularly high uncertainty.

In contrast, scaling overperformance is fairly sensitive to *β*, though only somewhat more so than to *P*_*tot*_ (and less than to *C*_*tot*_, which we can see from Figure 3 is still not particularly large (remember that overperformance is a *relative* measure of the effectiveness of CRISPRa vs CRISPRi). We are therefore less confident in our ability to quantitatively predict scaling overperformance than fold change overperformance, but we believe the overall trend that CRISPRa performs better than CRISPRi under conditions of high gRNA competition will hold.

